## Supplemental Tables 1-4 for "Longitudinal dynamics of the human vaginal ecosystem across the reproductive cycle"

**Supplementary Tables 1-4**

**Supplementary Table 1.** Demographic characteristics of study participants ($n$ = 178) enrolled at Stanford University (SU) and the University of Alabama at Birmingham (UAB)

| | SU | | UAB |
|---|---|---|---|
| Characteristic | Full cohort ($n$ = 82) | Postpartum subset ($n$ = 59) | Full cohort ($n$ = 96) |
| Median age at enrollment, years (range) | 32 (25 - 43) | 31 (25 - 43) | 25 (17 – 38) |
| Race: | | | |
| American Indian | 1 (1%) | 1 (2%) | 0 |
| Asian | 20 (24%) | 18 (31%) | 1 (1%) |
| Black | 4 (5%) | 4 (7%) | 80 (83%) |
| Other | 5 (6%) | 4 (7%) | 4 (4%) |
| Pacific Islander | 0 | 0 | 1 (1%) |
| White | 51 (62%) | 32 (54%) | 9 (9%) |
| Declined to state | 1 (1%) | 0 | 1 (1%) |
| Ethnicity: | | | |
| Hispanic | 13 (16%) | 5 (8%) | 5 (5%) |
| Non-Hispanic | 69 (84%) | 54 (92%) | 89 (93%) |
| Declined to state | 0 | 0 | 2 (2%) |
| Median pre-pregnancy BMI, $kg/m^2$ (range) | 22.5 (17.3 – 50.5) | 22.0 (17.3 – 50.5) | 28.0 (15.8 - 73.0) |

BMI, body mass index

**Supplementary Table 2.** Clinical characteristics of participants' pregnancies ($n$ = 196) enrolled at Stanford University (SU) and the University of Alabama at Birmingham (UAB)

| | SU | | UAB |
|---|---|---|---|
| Characteristic | Full cohort ($n$ = 100) | Postpartum subset ($n$ = 72) | Full cohort ($n$ = 96) |
| Nulliparous | 42 (42%) | 32 (44%) | 2 (2%) |
| History of prior live birth | 55 (55%) | 38 (53%) | 93 (97%) |
| History of prior preterm birth | 19 (19%) | 11 (15%) | 88 (92%) |
| Median birth-to-conception interval, months (range) | 25 (4 – 183) | 20 (5 – 183) | 31 (2 – 158) |
| Outcome: | | | |
| Miscarriage | 2 (2%) | 0 | 0 |
| Preterm delivery | 11 (11%) | 5 (7%) | 41 (43%) |
| Term delivery | 87 (87%) | 67 (93%) | 55 (57%) |
| Median gestational age at delivery, days (range) | 275 (219 – 293) | 275 (230 – 292) | 260 (123 – 289) |
| Median maternal age at delivery, years (range) | 33.0 (26.0 – 43.7) | 33.0 (26.0 – 43.7) | 26.5 (17.6 – 39.1) |
| Spontaneous labor onset | 39 (40%) | 29 (40%) | 66 (69%) |
| Spontaneous rupture of membranes (SROM) | 31 (32%) | 21 (29%) | 59 (61%) |
| GBS status, neg/pos/unk | 75/21/2 | 56/14/2 | 48/17/31 |
| Intrapartum antibiotics | 45 (46%) | 33 (46%) | 48 (50%) |
| Cesarean delivery | 30 (31%) | 19 (26%) | 22 (23%) |
| Female baby | 46 (47%) | 33 (46%) | 40 (42%) |

GBS, group B *Streptococcus*

**Supplementary Table 3.** Association between delivery and amplicon sequence variants (ASVs) in pairs of samples immediately flanking delivery (SU cohort, *n* = 70 pregnancies)

| **ASV** | **Change in (mean rank in) relative abundance** | **Effect size (*r*)** | **Significance code** |
|---|---|---|---|
| asv2 *Lactobacillus crispatus* | decreased | 0.63 | **** |
| asv4 *Lactobacillus jensenii* | decreased | 0.51 | **** |
| asv3 *Lactobacillus gasseri* | decreased | 0.42 | *** |
| asv1 *Lactobacillus iners* | decreased | 0.28 | * |
| asv11 *Ureaplasma parvum*/*urealyticum* | decreased | 0.27 | * |
| asv82 *Actinomyces neuii* | decreased | 0.08 | ns |
| asv25 *Staphylococcus aureus*/*epidermidis*/(others) | decreased | 0.05 | ns |
| asv43 *Prevotella buccalis* | increased | 0.75 | **** |
| asv21 *Peptoniphilus grossensis*/*vaginalis* | increased | 0.72 | **** |
| asv38 *Dialister propionicifaciens* | increased | 0.70 | **** |
| asv77 *Ezakiella coagulans* | increased | 0.69 | **** |
| asv26 *Anaerococcus mediterraneensis*/*murdochii* | increased | 0.67 | **** |
| asv115 *Peptoniphilus lacrimalis* | increased | 0.67 | **** |
| asv84 *Anaerococcus vaginalis* | increased | 0.66 | **** |
| asv133 *Peptoniphilus urinimassiliensis* | increased | 0.66 | **** |
| asv33 *Campylobacter ureolyticus* | increased | 0.65 | **** |
| asv100 *Clostridiales* KU726662 | increased | 0.65 | **** |
| asv14 *Prevotella timonensis* | increased | 0.64 | **** |
| asv64 *Peptoniphilus duerdenii* | increased | 0.64 | **** |
| asv102 *Porphyromonas asaccharolytica* | increased | 0.64 | **** |
| asv10 *Finegoldia magna* | increased | 0.64 | **** |
| asv88 *Porphyromonas uenonis* | increased | 0.62 | **** |
| asv59 *Peptoniphilus gorbachii*/*harei*/*lacydonensis*/*tyrrelliae* | increased | 0.61 | **** |
| asv106 *Mobiluncus curtisii* | increased | 0.60 | **** |
| asv81 *Peptoniphilus coxii* | increased | 0.59 | **** |
| asv36 *Lawsonella clevelandensis* | increased | 0.58 | **** |
| asv29 *Anaerococcus hydrogenalis* | increased | 0.57 | **** |
| asv60 *Varibaculum anthropi*/*cambriense* | increased | 0.57 | **** |
| asv87 *Campylobacter hominis* | increased | 0.56 | **** |
| asv47 *Fenollaria massiliensis* | increased | 0.56 | **** |
| asv93 *Porphyromonas bennonis* | increased | 0.53 | **** |
| asv39 *Anaerococcus obesiensis* | increased | 0.53 | **** |
| asv32 *Corynebacterium amycolatum*/*lactis*/*xerosis* | increased | 0.53 | **** |
| asv52 *Dialister micraerophilus* | increased | 0.52 | **** |
| asv56 *Anaerococcus prevotii*/*tetradius* | increased | 0.50 | *** |
| asv17 *Prevotella bivia* | increased | 0.48 | *** |
| asv54 *Fusobacterium nucleatum* | increased | 0.45 | *** |
| asv28 *Prevotella timonensis* | increased | 0.42 | *** |
| asv42 *Prevotella disiens* | increased | 0.38 | ** |
| asv31 *Corynebacterium tuberculostearicum* | increased | 0.34 | ** |
| asv20 *Streptococcus anginosus* | increased | 0.32 | ** |
| asv91 *Anaerococcus senegalensis* | increased | 0.27 | * |
| asv62 *Corynebacterium pseudogenitalium* | increased | 0.26 | * |
| asv78 *Corynebacterium pyruviciproducens* | increased | 0.23 | * |
| asv22 *Prevotella bivia* | increased | 0.23 | * |

Test performed if ASV had a relative abundance > 0.001 in at least 15% of tested samples (*n* = 45 ASVs; all shown). Significance levels correspond to Benjamini-Hochberg-adjusted *P*-values for paired Wilcoxon signed rank tests. ****, $P < 0.0001$; ***, $P < 0.001$; **, $P < 0.01$; *, $P < 0.05$; ns, not significant

**Supplementary Table 4.** Postpartum contraception methods

| Method | SU<br>Postpartum subset<br>(*n* = 72) |
|---|---|
| Birth control implant | 2 |
| Birth control injection | 1 |
| Birth control patch | 1 |
| Birth control pill | 3 |
| IUD, copper | 2 |
| IUD, progestin | 2 |
| IUD, unspecified type | 6 |
| Condoms | 2 |
| Tubal ligation | 1 |
| None | 12 |
| Did not respond | 40 |

IUD, Intrauterine device
